# Unified cell segmentation for spatial transcriptomics via transcript reassignment

**DOI:** 10.64898/2026.11.03.764126

**Authors:** Junkai Zhang, Xiyao Cheng, Juexin Wang, Wei Chen, Hao Chen

## Abstract

High-resolution spatial transcriptomics (ST) measures gene expression as individually localized molecules on imaging-based platforms or as counts in subcellular bins on sequencing-based platforms, and both must be assigned to cells before cell-level analysis. Existing segmentation methods focus on delineating cell boundaries, yet transcripts that diffuse beyond their cell of origin, fall where cells overlap, or lie in bins that cells only partially cover are still misassigned. Here we present Tracel, a two-stage optimal transport framework that assigns spatial observations to cells under the guidance of a matched single-cell RNA-sequencing reference. Tracel first aligns reference cells to spatial cells to estimate the expected expression profile at each cell location, and then maps transcripts to cells against these profiles. The same formulation assigns transcripts from imaging-based platforms (e.g., Xenium, CosMx) and bin counts from sequencing-based platforms (e.g., Visium HD). Across five human datasets spanning three platforms and three tissues, Tracel improves the separation of cell types over existing methods, assigning cell-type markers to their correct cells, retaining more cell-intrinsic transcripts and reducing contamination from neighboring cells. Tracel also preserves variation within cell types, yielding more accurate differential expression across spatial locations, and its expression profiles attribute ligand-receptor interactions to the correct cell-type pairs in cell-cell communication analysis. Tracel can enter the segmentation workflow at different stages, either refining an existing segmentation or segmenting cells starting from nucleus segmentation.

**Availability:** Code and example data are available at: https://tracel.readthedocs.io/en/latest/

## 1 Introduction

A cell’s identity is defined by the genes it expresses, but its function in a tissue also depends on its location and neighbors. Spatial transcriptomics (ST) captures both by measuring gene expression in its spatial context [1–3], with resolution that has advanced from multicellular spots to individual molecules. Imaging-based platforms such as MERFISH [4], Xenium [5] and CosMx [6] detect single mRNA molecules through cyclic in situ hybridization, recording the gene identity and coordinates of millions of transcripts. Sequencing-based platforms such as Visium HD [7] and Stereo-seq [8] capture transcripts on arrays of barcoded spots smaller than a cell (2 × 2 *µ*m bins for Visium HD; 0.5 *µ*m spacing for Stereo-seq).

At this resolution, however, measurements are no longer organized by cell. Each transcript must first be attributed to the cell that produced it, a step known as cell segmentation that underlies nearly all downstream analyses, from cell-type annotation and spatial organization to within-type expression variation and cell–cell communication. Transcripts merged from neighboring cells by segmentation errors produce spurious mixtures, leading to incorrect cell-type labels and biased expression estimates [9, 10].

Segmentation is commonly framed as delineating cell boundaries, yet what ultimately matters is which cell expressed each transcript, since this assignment defines the expression profiles used downstream. Most existing methods partition the tissue into non-overlapping regions, defined from nuclear or membrane stains (e.g., Cellpose, StarDist, Mesmer and CellSAM [11–14]) or with the help of transcripts (e.g., SCS and Proseg [15, 16]), and assign all transcripts within a region to its cell. Such a partition cannot account for common features of real tissue. Sections are typically 5-10 *µ*m thick [5, 7, 17], so cells overlap in the 2D projection. Diffusion during tissue processing displaces transcripts into neighboring cells. And on sequencing-based platforms, bins often only partially overlap a cell. Neither situation can be resolved by a mask-based assignment, regardless of how accurately the boundaries are drawn. Transcript-based methods such as Baysor and Segger instead assign individual transcripts, based on local gene composition or predicted transcript–nucleus links [9, 18], and can in principle handle these cases. However, they typically infer each cell’s expected composition from its observed transcripts, often those within the nucleus, and may miss cytoplasmic transcripts whose composition differs; they can also be slow on whole-slide data [18].

Here we present Tracel (**Tra**nscript to **cel**l mapping), which formulates cell segmentation as mapping each transcript to its cell of origin. This problem can be analogous to aligning sequencing reads to the reference genome in RNA-seq analysis. Unlike read alignment, however, segmentation lacks a known reference, the expression profile of each cell, and hence how many transcripts of each gene it should receive is unknown. We reasoned that this reference can be estimated from a matched single-cell RNA-seq (scRNA-seq) dataset.

Tracel proceeds in two optimal transport (OT) steps. In the cell OT step, reference cells are mapped onto initial spatial cells, which may come from nucleus segmentation, another segmentation method or the platform’s default pipeline (e.g., Xenium Onboard Analysis). We formulate this as a fused Gromov-Wasserstein OT problem that combines cross-dataset expression similarity with the similarity structure within each dataset, yielding an expected expression profile for each spatial cell. In the transcript OT step, unbalanced OT transports individual transcripts to nearby cells, with these profiles as the target distribution, using a cost defined by distance to candidate cells and constrained by gene identity. To accommodate extracellular transcripts, the transport is designed to be unbalanced, so that transcripts inconsistent with all nearby cells may remain unassigned. Each transcript is then assigned to the cell with the largest transport value. For sequencing-based data, counts in a bin are assigned to nearby cells gene by gene, so a single bin can contribute to several cells.

We applied Tracel to five human datasets spanning imaging-based (Xenium, CosMx) and sequencing-based (Visium HD) platforms and three tissues. Tracel improved cell-type separation over existing methods, assigning cell-type markers to their correct cells, retaining more cell-intrinsic transcripts and reducing contamination from neighboring cells. In addition, it shows similar performance when starting from an existing segmentation with or without cell-type annotation, making it practical to current ST pipelines. Although guided by a reference, Tracel preserves variation within cell types. In CosMx lung cancer tissue, it revealed that tumor cells near T cells showed a stronger interferon-*γ* response than distant tumor cells, while carrying less T cell marker expression than profiles from other methods, indicating a tumor-intrinsic difference rather than contamination. In cell-cell communication analysis, Tracel attributed ligand-receptor interactions to the correct cell-type pairs.

## 2 Methods

### 2.1 Tracel framework

Tracel assigns each observed transcript, or counts in a bin, to one of a predefined set of spatial cells in two optimal transport (OT) steps. The first step, cell OT (Fig. 1A), maps cells of a matched single-cell RNA-sequencing (scRNA-seq) reference (source) to spatial cells (target), with the cost between two cells defined by their expression dissimilarity in a joint PCA space. When cell-type annotations are available, a structure term in fused Gromov-Wasserstein (FGW) [19] encourages source cells sharing a label to be mapped to target cells sharing a label. Because labels are compared only within each domain, they do not need to have the same names or a one-to-one correspondence between the two domains. Cell OT yields an expected expression profile for each spatial cell, computed as the average expression of reference cells weighted by the transport plan. The second step (Fig. 1B), transcript OT, maps each observed transcript to nearby cells, using their expected expression of that gene as the target distribution. This is solved gene by gene and patch by patch, allowing parallel computation. Finally, the transport plan is binarized, assigning each transcript to a single cell to give an integer cell-by-gene matrix. Transcripts with low transport mass are treated as background and left unassigned. The expected profiles only guide the assignment and never enter the output. Every count is a transcript observed in the spatial data, and no value is imputed from the reference dataset.

**Figure 1.**
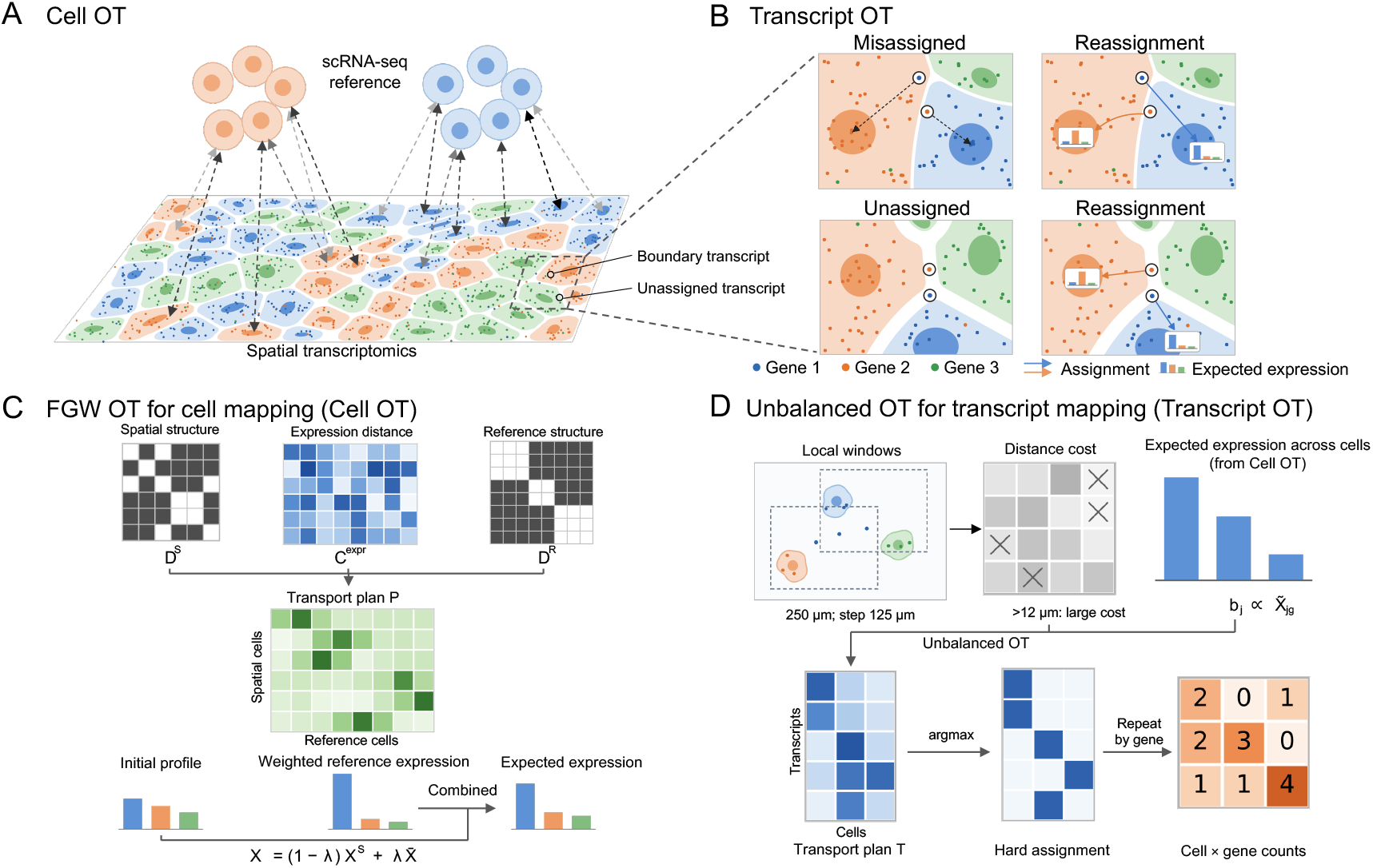
Overview of the Tracel framework. **A**, Cell OT maps cells of a tissue-matched scRNA-seq dataset to the initial spatial cells or nuclei. **B**, Enlarged view of a window in **A.** Transcript OT assigns each observation to a nearby cell according to the cells’ expected expression and distance, so an observation can be reassigned across initial cell segmentation boundaries. **C**, Cell OT, a fused Gromov–Wasserstein transport, with the objective that combines cross-dataset expression similarity with cell-group relationships within each dataset to estimate an expected expression profile for every spatial cell. **D**, Transcript OT, a gene-specific spatial transport, whose objective combines the expected expression profiles of cells with observation-to-cell distance. Each retained observation is assigned to one cell to produce an integer cell-by-gene matrix. In a Visium HD bin, one gene’s full UMI count contributes to one cell.

### 2.2 Input data format

The inputs are an initial spatial cell-by-gene count matrix, with the centroid of each cell as its location, a tissue-matched scRNA-seq count matrix, optional cell-type or cell-cluster labels for each, and spatial transcript records, each with a gene identity and coordinates. For Visium HD, each nonzero bin-gene pair is one record, located at the bin center and weighted by the UMI count of this gene. All UMIs of a gene in a bin are assigned together to a single cell. Genes were matched by symbol between two domains (spatial and single cell reference), and both matrices were restricted to the shared genes and normalized to 10^4^ counts per cell.

### 2.3 Reconstructing spatial cell profiles by mapping a single-cell reference

Cell OT requires two kinds of distance: an expression distance between cells of the two domains, and an optional structure distance between cells within each domain. Let 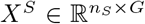 and 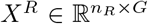 denote the normalized spatial and reference matrices over *G* shared genes. We concatenated the two matrices and applied a log(1 + *x*) transformation. Each gene was then z-scored, with values clipped at 10, and a principal component analysis (PCA) was performed on the concatenated matrix. The expression distance 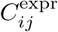 between spatial cell *i* and reference cell *j* is the squared Euclidean distance between their scores on the first 50 components of PCA. The structure distance is defined by cell-type labels. Each label is encoded as a one-hot vector with non-zero entry 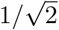, so that the squared distance between two cells of the same domain is 0 if they share a label and 1 otherwise. These distances form *D*^*S*^ and *D*^*R*^; cell-type labels are never compared across domains.

Cell OT (Fig. 1C) seeks a transport plan 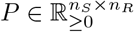, where *P*_*ij*_ is the mass transported between spatial cell *i* and reference cell *j*. Each cell carries a prescribed mass, its marginal (*a*_*i*_ for spatial and *b*_*j*_ for reference cells). *P* is found by minimizing the unbalanced fused Gromov–Wasserstein (FGW) objective, which keeps the row and column sums of *P* close to these marginals:

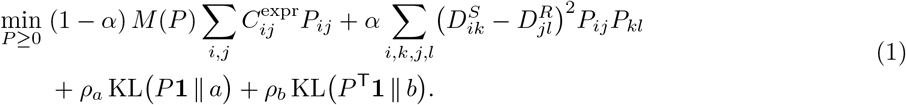

The marginals are type-balanced: a spatial cell in a group of 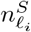 cells among *K*_*S*_ groups receives 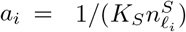, and likewise 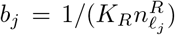 for reference cells, so that large cell types do not dominate the transport. *M* (*P*) = ∑_*ij*_*P*_*ij*_ is the total transported mass, and *α* weights the structure term against the expression term. The Kullback–Leibler (KL) terms penalize deviations of the row and column sums of *P* from *a* and *b* rather than enforcing equality, so that cells without a good counterpart in the other domain can transport less mass. The structure term can be omitted when cell-type labels are unavailable. We solved Eq. 1 with the low-rank FGW solver of moscot [20, 21], which factorizes the plan as *P* = *Q* diag(*g*)^−1^*R*^T^ for efficient computation on large datasets, using rank 20, *α* = 0.9, marginal relaxation parameters *τ*_*a*_ = 0.8 and *τ*_*b*_ = 0.9 (*τ* = 1 gives balanced transport).

For each spatial cell, we computed its transported mass *ρ*_*i*_ = ∑_*j*_*P*_*ij*_ and the average reference expression weighted by the transport plan, 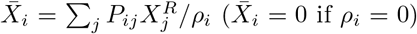. A cell that transports less than *a*_*i*_ is poorly matched in the reference, so its expected expression profile 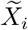 combines 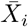 with the cell’s own expression, weighted by the fraction of its mass transported:

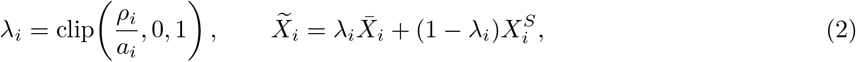

so the expected expression profile is obtained in the normalized count space.

### 2.4 Mapping transcripts to cells guided by reconstructed profiles

Transcript OT (Fig. 1D) was solved separately for each gene within each window (next section). Let ℳ be the records of gene *g* in window *W*. The cells within 12 *µ*m of at least one record in ℳ, found with a KD-tree in SciPy [22], form the candidate cell collection J in the window *W*. For record *m* and candidate cell *j* at distance *d*_*mj*_ (*µ*m), the cost was:

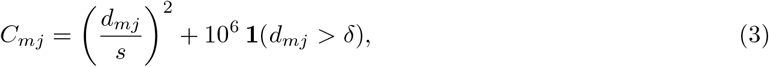

where *s* is set as 15, which is a distance scale that controls how quickly the attraction of a cell for a transcript decays with distance. In the optimal plan of the entropically regularized problem (below), the attraction of a cell for a transcript decays with distance as 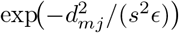, where *ϵ* is the entropic regularization weight. The second term excludes pairs farther than *δ* apart, which is set as 12 *µm* across all the five datasets we studied. The record-side marginal *a*_*m*_ was proportional to the record weight *w*_*m*_ (*w*_*m*_ = 1 for measured transcripts and the UMI count for Visium HD), normalized by the total counts of *g* in window *W*. The cell-side marginal *b*_*j*_ was proportional to *X*_*jg*_, normalized over J, so that the target share of each cell is proportional to its expected expression of *g*. The transport plan *T* solved the entropically regularized unbalanced OT problem [23, 24], using the Sinkhorn solver of OTT-JAX [25]:

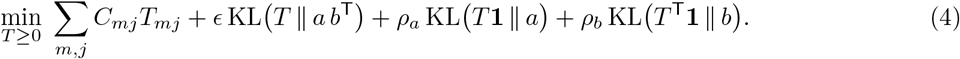

The marginal terms allow a record to transport less than *a*_*m*_, so that records inconsistent with all nearby cells can remain unassigned. We used *ϵ* = 0.05, *τ*_*a*_ = 0.8 and *τ*_*b*_ = 0.9, where *τ* = *ρ/*(*ρ* + *ϵ*) is the marginal relaxation parameter in the range of (0, 1].

### 2.5 Sliding-window computation of transcript OT

To keep each problem small, we tiled the tissue with 250 × 250 *µ*m windows placed every 125 *µ*m in *x* and *y*, so that adjacent windows overlap by half their width and every location away from the border is covered by four windows. Records were partitioned by window, whereas candidate cells were searched over all cells, so a record near a window edge can still be assigned to a cell outside the window. Windows are independent and were processed in parallel.

Within each window, a record was assigned to *z*_*m*_ = arg max_*j*_*T*_*mj*_*/* ∑_*k*_*T*_*mk*_, and this maximum share was kept as its assignment confidence. Because of the overlap, each record is solved in several windows, at least one of which contains it away from the edge; we kept the assignment with the highest confidence. This assignment was retained only if:

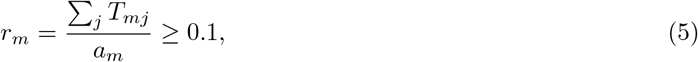

with *a*_*m*_ the record’s marginal in that window; otherwise the record was left unassigned. The final count matrix was:

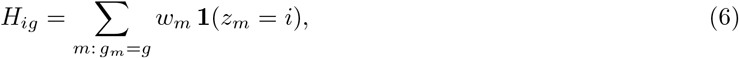

summed over retained assignments. Each measured transcript contributes one count, and each Visium HD record contributes its full UMI count to one cell. *H* is integer-valued and contains only observed transcripts.

### 2.6 Datasets and preprocessing

We applied Tracel to five human cancer datasets from Xenium, CosMx and Visium HD, covering breast, lung and colon, each paired with a published scRNA-seq reference of the same tissue and cancer type. Xenium breast S1 (10x Genomics FFPE breast cancer “Sample 2” [5, 26], 288-gene panel, 142,272 cells in initial segmentation) used the cell-type annotations released with the dataset, derived from unsupervised clustering, with the breast cancer atlas of Wu et al. [27] (SCP1039; 100,064 cells) as reference. Xenium breast S2 (replicate 1 of Janesick et al. [5], 313-gene panel, 167,780 cells) used the study’s cell-type annotations, which were transferred from the matched scFFPE-seq data (27,472 cells) used here as the reference. CosMx lung (NanoString NSCLC Lung5 Rep2 [6], 960-gene panel, 105,800 cells) and Xenium lung (10x Genomics lung cancer, 377-gene panel [28], a contiguous region of 49,786 cells are selected for study) both used tumor-lung cells of the lung adenocarcinoma atlas GSE131907 [29] as reference (45,149 cells). For CosMx lung, the official annotations released by NanoString with the dataset were mapped to eight broad groups to match the resolution of the reference (Supplementary Note 1 and Table S1). For Xenium lung, which has no cell-type annotation, we annotated the Xenium graph-based clusters by marker genes and mapped them to seven broad groups (Supplementary Tables S2 and S3). Visium HD colon is sample P2 of Oliveira et al. [7], with counts in 2 *µ*m bins. We detected 49,927 nuclei in a 2.7 × 2.7 mm region with a pretrained StarDist H&E model [12] and assigned each bin to the nearest nucleus centroid within 12 *µ*m to form the initial cells. Each cell took the published deconvolution label of the nearest 8 *µ*m bin, and the scRNA-seq reference (35,298 cells) came from the same study. Because the platform measures the full transcriptome, cell OT used all shared genes, whereas, to reduce computation, transcript OT was run only on the 2,000 genes whose expected profiles differed most from the initial profiles (mean absolute difference across cells). Evaluation used the full transcriptome. For the four imaging-based datasets, the initial cell segmentation came from each platform’s own pipeline, which segments cells from the paired DAPI or membrane protein images.

## 3 Results

### 3.1 Tracel improves cell-type separation across spatial transcriptomics platforms

In spatial transcriptomics analyses, cell types are commonly less clearly separated than in scRNA-seq data, partly because segmentation errors mix transcripts from neighboring cells [10, 16, 30]. We therefore first assessed how well different methods separate cell types. We compared Tracel with the initial segmentation from which it started (Original), the image-based segmentation method Cellpose [11, 31], and the transcript-based methods Proseg [16] and Baysor [9]. Cellpose segmented nuclei from the DAPI image, and transcripts were assigned either only to the nucleus containing them (Cellpose nucleus) or, if outside all nuclei, also to the nearest nucleus within 12 *µ*m as an approximation of the whole cell (Cellpose cell). Within each dataset, all methods were evaluated on the same gene set. We first examined whether the expression profiles from each method separate distinct cell populations, by visualizing them in UMAP [32] embeddings colored by the published cell-type annotations (Fig. 2A). Across all five datasets, Tracel produced tighter clusters with clearer separation between the clusters than Original and the other segmentation methods. For example, in Xenium breast S1 (the first row in Fig. 2A), B cells were intermixed with *TRAC* ^+^ T cells in Original and the other methods but formed a separate cluster with Tracel. And even clearer in Visium HD colon (the last row in Fig. 2A), Myofibroblasts were scattered among plasma, goblet and endothelial cells by all other methods but formed a distinct cluster with Tracel.

**Figure 2.**
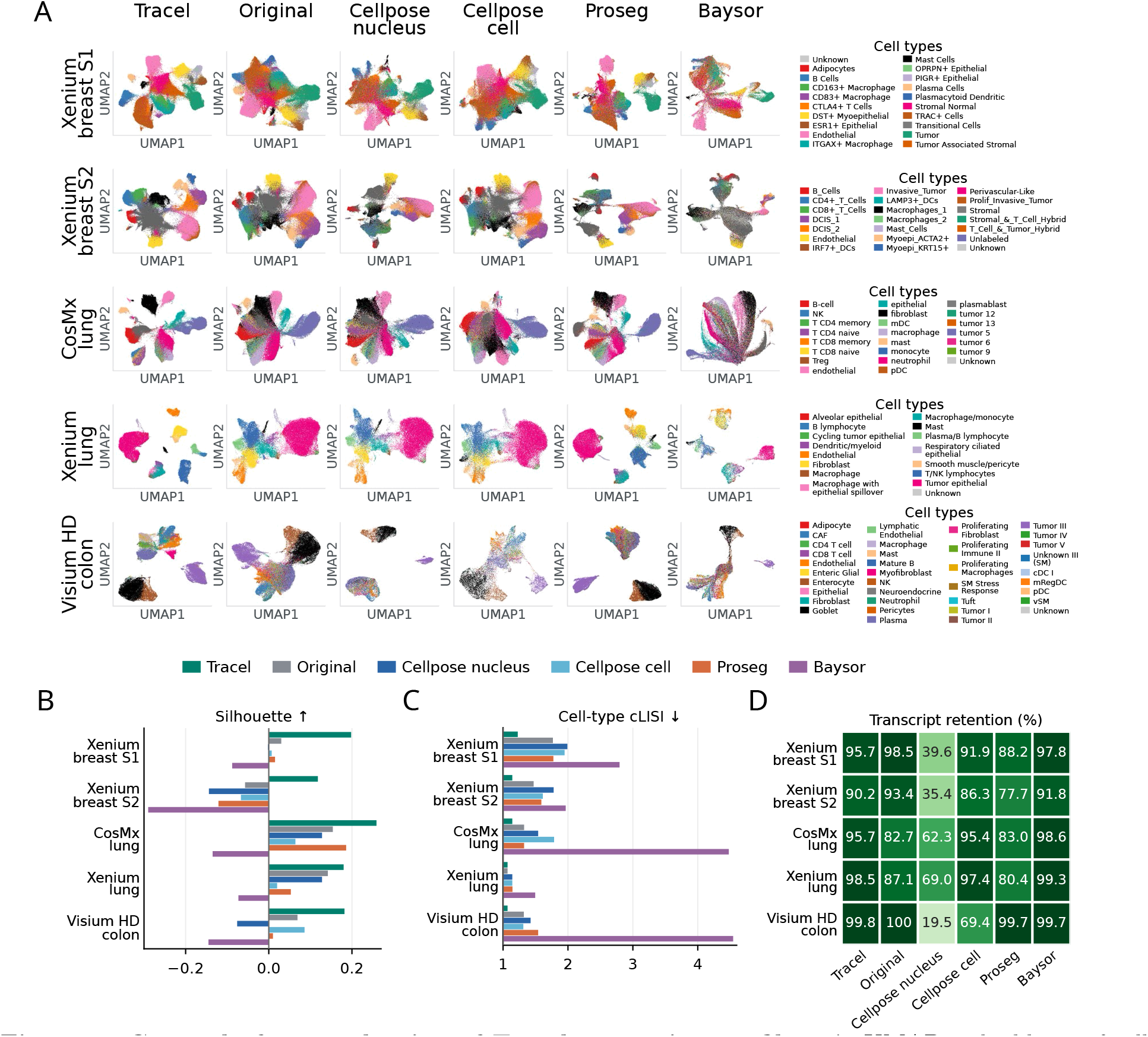
Cross-platform evaluation of Tracel expression profiles. **A**, UMAP embeddings of cells based on their expression profiles produced by Tracel, the Original segmentation, Cellpose nucleus, Cellpose cell, Proseg and Baysor across the five datasets. Cells are colored by their published cell-type annotations. **B**, Mean silhouette scores of different methods across five datasets. Higher scores indicate stronger separation of labeled cell types in UMAP. **C**, Median cell-type LISI (cLISI), calculated from 30 nearest neighbors in the PCA embedding of cells produced by each method. Lower values indicate less local mixing of cell-type labels with the minimum value of 1. **D**, Percentage of observed transcripts or UMI counts retained in the final cell-by-gene matrices. For Visium HD colon, retention was calculated only for the 2,000 genes reassigned by transcript OT, whereas in **A**–**C** evaluation in the Visium HD colon dataset uses all 18,079 genes.

We then quantified this separation with two label-based metrics: the mean silhouette score [33], which measures how well cells of each type are separated from cells of other types (higher is better), and the median cell-type LISI (cLISI) [34], which measures the local mixing of cell types (lower is better). Because the silhouette score relies on Euclidean distances between all cells, which in PCA space may not follow the nonlinear structure of the data, we computed it on each method’s two-dimensional UMAP embedding, which is designed to preserve local manifold structure. cLISI depends only on local neighborhoods and was computed from the 30 nearest neighbors in each method’s 30-component PCA embedding. Because true cell identities are unavailable, these metrics rely on the published annotations, which were derived from the Original segmentation and thus favor Original, and which Tracel also used in its cell OT step. Original therefore is the primary comparison. The other methods produce different cells, which took the labels of the annotated cells: Proseg cells kept the label of the Original cell they started from, and Cellpose and Baysor cells took the label of the nearest Original cell centroid within 20 *µ*m (cells without such a match were not scored). Numbers for these methods serve as references.

Tracel achieved the highest silhouette score in all five datasets (Fig. 2B), increasing it relative to Original from 0.030 to 0.198 in Xenium breast S1, from −0.057 to 0.118 in Xenium breast S2, from 0.154 to 0.259 in CosMx lung, from 0.142 to 0.180 in Xenium lung, and from 0.069 to 0.182 in Visium HD colon. It also achieved the lowest cLISI in all five datasets (Fig. 2C; 1.23, 1.14, 1.14, 1.07 and 1.07, versus 1.77, 1.47, 1.32, 1.07 and 1.31 for the best other method). Notably, in CosMx lung and Xenium lung, Tracel separated the finer cell types shown in Fig. 2A even though cell OT used only broad groups.

Tracel remains effective when cell-type labels are unavailable. We evaluated a variant without the cell-type structure term in cell OT (Tracel no-structure), and the same variant initialized from the nucleus segmentation (Tracel nucleus-only). Both variants still produced more separated cell-type clusters compared to Original (Supplementary Fig. S1A). And even without cell-type labels, Tracel no-structure achieved the best silhouette and cLISI scores among all compared methods (Supplementary Fig. S1B-C).

Besides achieving better separation of cell types, Tracel also retained most transcripts (95.70%, 90.15%, 95.65%, 98.53% and 99.84% in Xenium breast S1, Xenium breast S2, CosMx lung, Xenium lung and Visium HD colon, respectively), comparable to Baysor and Original and higher than Proseg and Cellpose (Fig. 2D). Proseg (77.68–88.18% in the four imaging-based datasets) and Cellpose nucleus (19.50–69.00%) discarded many transcripts, possibly because they capture only nuclear regions or only transcripts resembling those of the nucleus from which they start, and thus recover only part of each cell’s profile. In contrast, Tracel reassigned most transcripts to cells, recovering more complete cell profiles.

### 3.2 Tracel assigns transcripts to their cells of origin

We next asked whether this improved separation arises because transcripts are assigned to their cells of origin. We first compared cell-type-specific expression between the spatial and scRNA-seq data: for each cell type, we computed one-versus-rest log_2_ fold changes in both datasets and measured their Pearson correlation. Tracel achieved the highest mean correlation between matching cell types among all methods in all five datasets (Fig. 3A; 0.801, 0.779, 0.738, 0.729 and 0.383), exceeding Original by 0.050, 0.140, 0.159, 0.083 and 0.010 in Xenium breast S1, Xenium breast S2, CosMx lung, Xenium lung and Visium HD colon, respectively. Thus, the cell-type-specific genes of Tracel profiles agree better with the reference.

**Figure 3.**
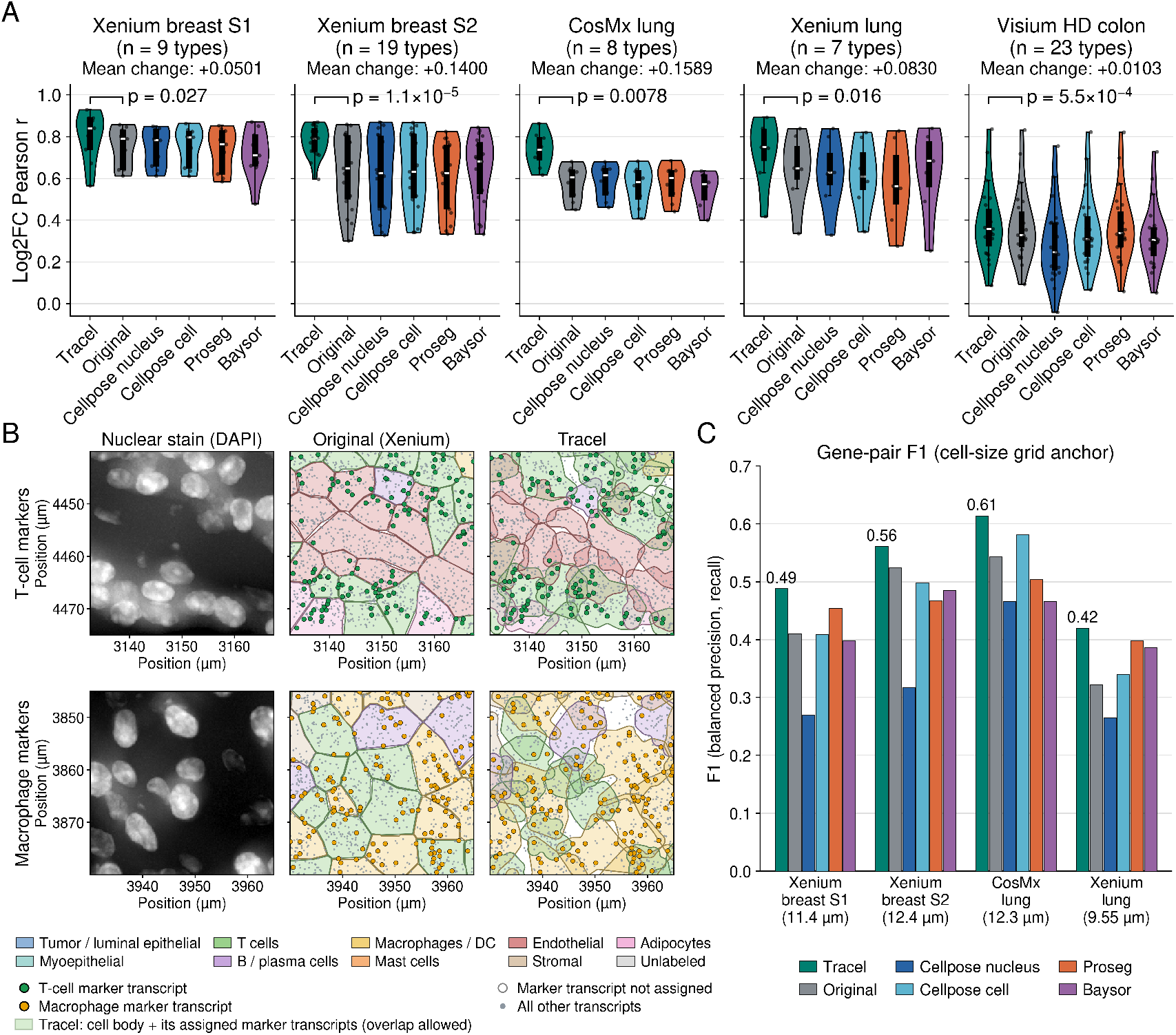
Tracel assigns transcripts to their cells of origin. **A**, Agreement between spatial and scRNA-seq cell-type expression signatures. For each cell type, we computed a one-versus-rest log_2_ fold-change profile from the spatial data and from the matched scRNA-seq reference, and calculated the Pearson correlation between the two. Each point is one cell type; distributions are shown for six methods in five datasets. Within each dataset, all methods are evaluated on the same set of cell types. Mean change is the mean correlation of Tracel minus that of Original; *P* values are from two-sided paired Wilcoxon signed-rank tests comparing Tracel with Original. **B**, Example regions in Xenium breast S1 showing T-cell markers (top) and macrophage markers (bottom). Left, nuclear stain; middle, Original cell boundaries; right, Tracel cell regions.In the Original segmentation, many T-cell and macrophage marker transcripts fall within the boundaries of cells of other types, whereas Tracel assigns them to cells of the corresponding type. **C**, Gene-pair F1 scores in the four imaging-based datasets. Grid sizes (in parentheses) are set relative to the median cell diameter in each dataset.

We then located these reassignments in the tissue. In Original, marker transcripts near cell boundaries were often assigned to adjacent cells of another type, whereas Tracel assigned them to cells of the expressing type (Fig. 3B). In a region of Xenium breast S1 where T cells border endothelial cells, only 40% of T-cell marker transcripts (*CD3E, CD3G, TRAC, CD247* and *CD8A*) were counted in T cells in Original, with many falling in neighboring endothelial cells; Tracel raised this fraction to 88% (Fig. 3B, top). Likewise, in a region where macrophages border T cells, the fraction of macrophage marker transcripts (*CD68, LYZ, CD163, C1QA, C1QC, CD14* and *MRC1*) counted in macrophages increased from 45% to 91% (Fig. 3B, bottom).

Finally, to test beyond marker genes whether each method’s profiles reflect genes expressed by the cell itself rather than a mixture with its neighbors, we used gene pairs identified from the Original segmentation (Supplementary Note 2): “intrinsic pairs”, enriched between a cell’s nucleus and its own cytoplasm, and “suspicious pairs”, enriched between a nucleus and the cytoplasm of an adjacent cell of a different type. We summarized purity (few suspicious pairs) and completeness (many intrinsic pairs recovered) in a single F1 score. Visium HD colon was excluded, because a bin can mix nuclear and cytoplasmic transcripts and span parts of different cells. Tracel achieved the highest F1 in all four imaging-based datasets (0.49, 0.56, 0.61 and 0.42 for Xenium breast S1, Xenium breast S2, CosMx lung and Xenium lung, versus at most 0.45, 0.52, 0.58 and 0.40 for the other methods; Fig. 3C), producing expression profiles that are both pure and complete.

### 3.3 Tracel retains variations within cell types across spatial locations

Cells of the same type can adopt different states depending on their surroundings, and differential expression between locations within a cell type is a common way to study these states. For example, tumor cells exposed to interferon-*γ* (IFN-*γ*) secreted by nearby T cells up-regulate MHC class I antigen presentation [37, 38], which makes them visible to cytotoxic T cells and is associated with response to immunotherapy [36, 39, 40]. Detecting such differences in spatial data is complicated by segmentation errors: tumor cells adjacent to T cells receive T-cell transcripts from their neighbors, so a difference between tumor cells near and far from T cells may reflect contamination rather than tumor-intrinsic biology. Conversely, because Tracel is guided by a scRNA-seq reference, it could in principle make cells of the same type more similar to each other and remove genuine within-type variation. We therefore tested whether Tracel removes contamination while preserving location-dependent expression differences within a cell type.

We compared tumor cells in the CosMx lung dataset that were close to T cells (at least three T cells within 50 *µ*m; *n* = 625) with tumor cells far from T cells (nearest T cell more than 100 *µ*m away; *n* = 7,857) (Fig. 4A; Supplementary Note 3). In Original, most T-cell markers were elevated in tumor cells adjacent to T cells (log_2_ fold change up to 0.7), indicating contamination from neighboring T cells; after Tracel, these fold changes were close to zero (Supplementary Fig. S2). In contrast, the antigen-presentation machinery (APM) module was significantly enriched by GSEA only after Tracel (normalized enrichment score (NES) 1.48, FDR = 0.36 in Original; NES 1.86, FDR = 0.003 in Tracel, Fig. 4B), and four of the six APM genes were significantly up-regulated after Tracel, compared with three in Original (Supplementary Fig. S2). The broader Grasso IFN-*γ* gene set showed the same trend, with NES increasing from −0.99 in Original to 1.16 in Tracel (Fig. 4C). As a function of distance to the nearest T cell, APM expression in tumor cells increased toward T cells over the first 50 *µ*m (Fig. 4D). In Original, however, it dropped within 15 *µ*m of a T cell, likely because contaminating T-cell transcripts take up a share of the normalized counts. IFN-*γ*-induced genes showed the same pattern (Fig. 4E). Consistently, contamination in Original was strongest within 15 *µ*m of T cells, whereas in Tracel it remained low at all distances (Fig. 4F).

**Figure 4.**
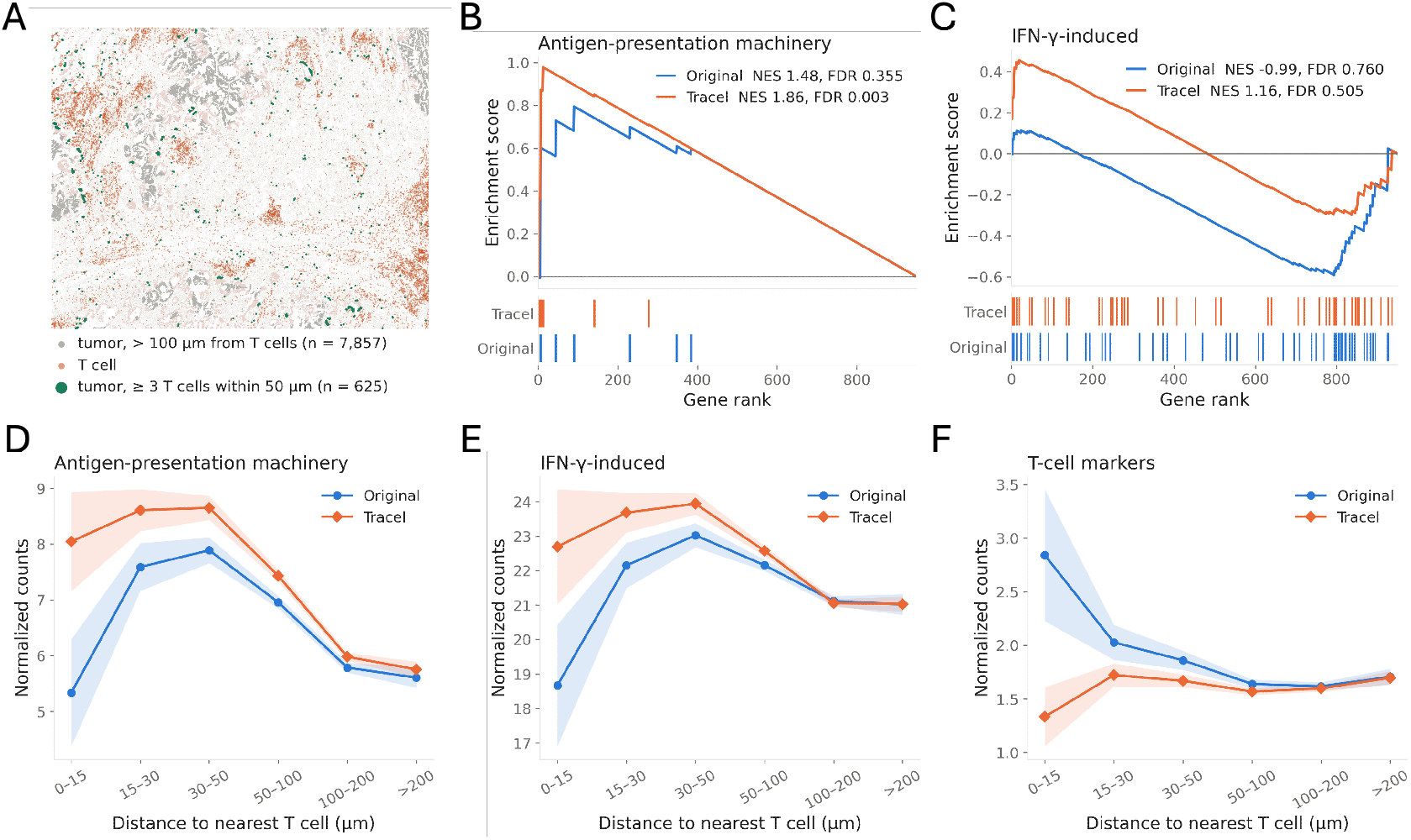
Tracel removes T-cell contamination from tumor cells while preserving their IFN-*γ* response in the CosMx lung dataset. **A**, Locations of tumor cells close to T cells (at least three T cells within 50 *µ*m; *n* = 625, green), tumor cells far from T cells (nearest T cell more than 100 *µ*m away; *n* = 7,857, gray), and T cells (orange). **B**,**C**, Gene set enrichment analysis of the differential expression between near and far tumor cells for the MHC class I antigen-presentation machinery (APM) gene set [35] (**B**) and the IFN-*γ*-induced gene set [36] (**C**), for Original (blue) and Tracel (orange). Tick marks below show the ranks of gene-set members, and the legends give the normalized enrichment score (NES) and false discovery rate (FDR). **D**–**F**, Expression of APM genes (**D**), IFN-*γ*-induced genes (**E**) and T-cell markers (**F**) in tumor cells, binned by the distance to the nearest T cell.

Together, these results show that Tracel removes transcripts contaminating tumor cells from adjacent T cells while preserving, and sharpening, the tumor-intrinsic IFN-*γ* response. Although guided by a reference, Tracel retains expression variation within a cell type across spatial locations, enabling differential expression analysis between cell states.

### 3.4 Tracel improves inference of cell–cell communication

We further tested whether Tracel’s expression profiles improve cell–cell communication analysis. We applied SpatialCellChat [41, 42] to the Original and Tracel expression profiles of the same 18,916 cells in the Xenium breast S1 dataset, using identical cell-type labels and analysis settings. Tracel reshaped the inferred communication network among cell populations (Fig. 5A). At the ligand–receptor level, 18 significant communications were shared, whereas 34 were significant only in Original and 33 only in Tracel. At the pathway level, 18 communications were shared, with 31 significant only in Original and 30 only in Tracel (Fig. 5B). Fewer interactions between different cell types were detected with Tracel, likely because in Original, transcripts misassigned between adjacent cells of different types create spurious interactions.

**Figure 5.**
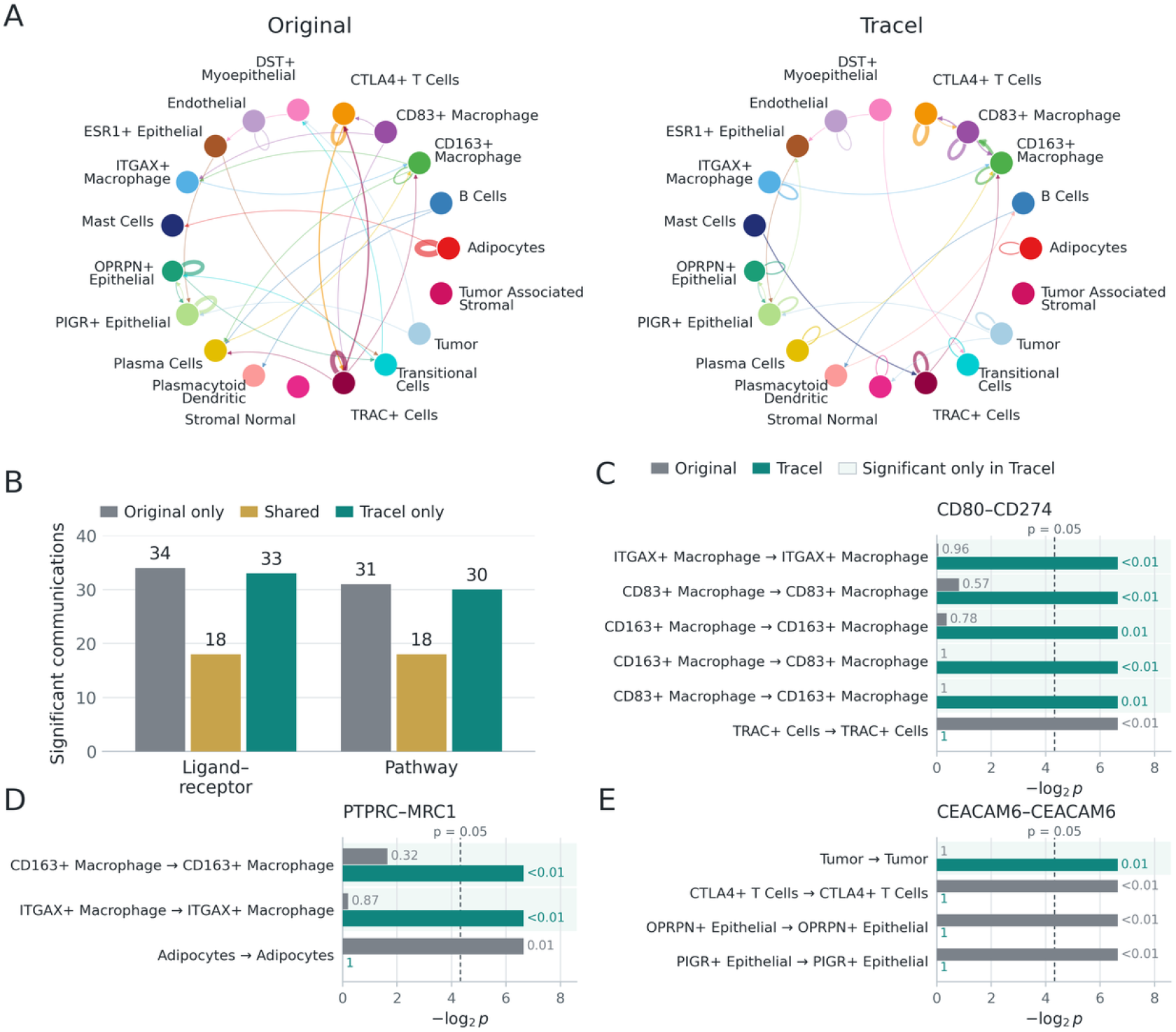
Tracel refines inferred cell–cell communication in the Xenium breast S1 dataset. **A**, Aggregated cell–cell communication networks inferred by SpatialCellChat from the Original and Tracel expression profiles. Both analyses use the same 18,916 cells, the same cell-type labels and the same settings. Edge color indicates the sender cell type, and edge width is proportional to the aggregated interaction weight, drawn on a scale shared by both methods. **B**, Numbers of significant ligand–receptor interactions and signaling pathways detected only in Original, in both methods, or only in Tracel. **C**–**E**, Significance of the CD80–CD274 (**C**), PTPRC–MRC1 (**D**) and CEACAM6–CEACAM6 (**E**) interactions for each sender–receiver pair that is significant in at least one method. Bars show −log_2_ *P* from the Spatial-CellChat permutation test (100 permutations) for Original (gray) and Tracel (cyan), with *P* values labeled. When no permutation reached the observed score (*P <* 0.01), the bar is drawn at *P* = 0.01. The dashed line marks *P* = 0.05; shading marks pairs significant only in Tracel.

Selected ligand–receptor signals changed in their inferred cellular context (Fig. 5C–E). CD80–CD274 became significant within *CD163* ^+^, *CD83* ^+^ and *ITGAX* ^+^ macrophages and between *CD163* ^+^ and *CD83* ^+^ macrophages after Tracel, whereas its signal within *TRAC* ^+^ T cells was no longer significant (Fig. 5C). PTPRC–MRC1 became significant within *CD163* ^+^ and *ITGAX* ^+^ macrophages, while its signal within adipocytes lost significance (Fig. 5D). For CEACAM6–CEACAM6, the signal within tumor cells became significant, whereas the signals within *CTLA4* ^+^ T cells and within *OPRPN* ^+^ and *PIGR*^+^ epithelial cells were no longer significant (Fig. 5E).

These changes placed CD80–CD274 and PTPRC–MRC1 signaling in macrophages, consistent with the expression of these molecules in antigen-presenting cells, and removed CEACAM6 signaling from T cells, where its expression is not expected. Together, these results suggest that Tracel improves downstream analysis of spatial transcriptomics data.

## 4 Discussion

Tracel treats cell segmentation as the assignment of individual spatial measurements to cells, guided by the expression expected for each cell from a matched scRNA-seq reference. Across five datasets from three platforms, it improved cell-type separation while retaining most transcripts, and its profiles changed downstream differential expression and cell–cell communication results.

Tracel differs from transcript decontamination. Decontamination methods developed for spatial data [10, 43–45] start from cell types and remove from each cell the transcripts that are unlikely to belong to its type. Tracel instead starts from an expected expression profile for each individual cell and assigns transcripts to reconstruct that cell’s own profile. It is therefore a segmentation of single cells rather than a cleanup at the level of cell types, and it preserves variation among cells of the same type. This difference also makes Tracel more flexible, it can start from an existing segmentation or from nuclei alone, and can work without cell-type labels (Supplementary Fig. S1).

The cell-type level evaluation is limited by the lack of ground truth, since the cell of origin of each transcript is unknown in real tissue. We therefore relied on published cell-type annotations, which are the only cell-level information available across datasets. These annotations were derived from the original segmentation and inherit its errors, and when used in cell OT they also guide Tracel itself. The most informative comparison is therefore with the original segmentation, against which Tracel recovered expression profiles that better separated cell types. Simulations with known transcript origin, membrane stains, or serial sections would provide more direct benchmarks.

Several extensions could further improve Tracel. Because it preserves the initial cell set, missing or merged cells cannot be resolved by transcript reassignment alone, and joint refinement of cells and transcript assignments could address these cases. The expected profiles depend on the quality and coverage of the reference. Cells without a good reference match fall back partly to their own expression, but rare populations or disease-associated states absent from the reference may be assigned less accurately. Cell OT is currently solved with moscot [20]. Other OT solvers and formulations could also be explored, such as total-variation-relaxed fused partial Gromov–Wasserstein, as in SpaOT [46]. For Visium HD, each bin-gene record is currently assigned as a whole. Dividing UMI counts among candidate cells may further improve reconstruction near cell boundaries.

Tracel is implemented in Python and is publicly available at https://tracel.readthedocs.io/en/latest/ including tutorials on the datasets used in this study. It provides a common framework for reconstructing cell expression profiles from imaging- and sequencing-based spatial transcriptomics, as a step before downstream analyses at the cell level.

## Notes

https://tracel.readthedocs.io/en/latest/

### Competing Interest Statement

The authors have declared no competing interest.

## References

[1] Patrik L Ståhl et al. “Visualization and analysis of gene expression in tissue sections by spatial transcriptomics”. In: Science 353.6294 (2016), pp. 78–82. DOI: 10.1126/science.aaf2403.

[2] Anjali Rao et al. “Exploring tissue architecture using spatial transcriptomics”. In: Nature 596.7871 (2021), pp. 211–220. DOI: 10.1038/s41586-021-03634-9.

[3] Lambda Moses and Lior Pachter. “Museum of spatial transcriptomics”. In: Nature Methods 19.5 (2022), pp. 534–546. DOI: 10.1038/s41592-022-01409-2.

[4] Kok Hao Chen et al. “Spatially resolved, highly multiplexed RNA profiling in single cells”. In: Science 348.6233 (2015), aaa6090. DOI: 10.1126/science.aaa6090.

[5] Amanda Janesick et al. “High resolution mapping of the tumor microenvironment using integrated single-cell, spatial and in situ analysis”. In: Nature Communications 14.1 (2023), p. 8353. DOI: 10.1038/s41467-023-43458-x. URL: https://www.nature.com/articles/s41467-023-43458-x.

[6] Shanshan He, Ruchir Bhatt, Carl Brown, et al. “High-plex imaging of RNA and proteins at subcellular resolution in fixed tissue by spatial molecular imaging”. In: Nature Biotechnology 40.12 (2022), pp. 1794–1806. DOI: 10.1038/s41587-022-01483-z. URL: https://www.nature.com/articles/s41587-022-01483-z.

[7] Michelli Faria de Oliveira et al. “High-definition spatial transcriptomic profiling of immune cell populations in colorectal cancer”. In: Nature Genetics 57.6 (2025), pp. 1512–1523. DOI: 10.1038/s41588-025-02193-3. URL: https://www.nature.com/articles/s41588-025-02193-3.

[8] Ao Chen et al. “Spatiotemporal transcriptomic atlas of mouse organogenesis using DNA nanoball-patterned arrays”. In: Cell 185.10 (2022), 1777–1792.e21. DOI: 10.1016/j.cell.2022.04.003.

[9] Viktor Petukhov et al. “Cell segmentation in imaging-based spatial transcriptomics”. In: Nature Biotechnology 40.3 (2022). Published online 14 October 2021, pp. 345–354. DOI: 10.1038/s41587-021-01044-w. URL: https://www.nature.com/articles/s41587-021-01044-w.

[10] Jonathan Mitchel et al. “Impact and correction of segmentation errors in spatial transcriptomics”. In: Nature Genetics 58.2 (2026), pp. 434–444. DOI: 10.1038/s41588-025-02497-4. URL: https://www.nature.com/articles/s41588-025-02497-4.

[11] Carsen Stringer et al. “Cellpose: a generalist algorithm for cellular segmentation”. In: Nature methods 18.1 (2021), pp. 100–106. DOI: 10.1038/s41592-020-01018-x.

[12] Uwe Schmidt et al. “Cell detection with star-convex polygons”. In: International Conference on Medical Image Computing and Computer-Assisted Intervention. Springer. 2018, pp. 265–273. DOI: 10.1007/978-3-030-00934-2_30.

[13] Noah F. Greenwald et al. “Whole-cell segmentation of tissue images with human-level performance using large-scale data annotation and deep learning”. In: Nature Biotechnology 40.4 (2022). Published online 18 November 2021, pp. 555–565. DOI: 10.1038/s41587-021-01094-0. URL: https://www.nature.com/articles/s41587-021-01094-0.

[14] Markus Marks et al. “CellSAM: a foundation model for cell segmentation”. In: Nature Methods 22.12 (2025), pp. 2585–2593. DOI: 10.1038/s41592-025-02879-w. URL: https://www.nature.com/articles/s41592-025-02879-w.

[15] Hao Chen, Dongshunyi Li, and Ziv Bar-Joseph. “SCS: cell segmentation for high-resolution spatial transcriptomics”. In: Nature Methods 20.8 (2023), pp. 1237–1243. DOI: 10.1038/s41592-023-01939-3. URL: https://www.nature.com/articles/s41592-023-01939-3.

[16] Daniel C. Jones et al. “Cell simulation as cell segmentation”. In: Nature Methods 22.6 (2025), pp. 1331–1342. DOI: 10.1038/s41592-025-02697-0. URL: https://www.nature.com/articles/s41592-025-02697-0.

[17] Jeffrey R. Moffitt et al. “Molecular, spatial, and functional single-cell profiling of the hypothalamic preoptic region”. In: Science 362.6416 (2018), eaau5324. DOI: 10.1126/science.aau5324.

[18] Elyas Heidari, Andrew Moorman, et al. “Segger: Fast and accurate cell segmentation of imaging-based spatial transcriptomics data”. In: bioRxiv (2025). Preprint. DOI: 10.1101/2025.03.14.643160. URL: https://www.biorxiv.org/content/10.1101/2025.03.14.643160v1.

[19] Titouan Vayer et al. “Optimal Transport for structured data with application on graphs”. In: Proceedings of the 36th International Conference on Machine Learning. Vol. 97. Proceedings of Machine Learning Research. PMLR, 2019, pp. 6275–6284. URL: https://proceedings.mlr.press/v97/titouan19a.html.

[20] Dominik Klein et al. “Mapping cells through time and space with moscot”. In: Nature 638.8052 (2025), pp. 1065–1075. DOI: 10.1038/s41586-024-08453-2. URL: https://www.nature.com/articles/s41586-024-08453-2.

[21] Meyer Scetbon et al. “Unbalanced Low-rank Optimal Transport Solvers”. In: Advances in Neural Information Processing Systems. Vol. 36. 2023, pp. 52312–52325. DOI: 10.52202/075280-2277. URL: https://proceedings.neurips.cc/paper_files/paper/2023/hash/a439259e78294c38d157a51a2c40486b-Abstract-Conference.html.

[22] Pauli Virtanen, Ralf Gommers, Travis E. Oliphant, et al. “SciPy 1.0: fundamental algorithms for scientific computing in Python”. In: Nature Methods 17.3 (2020), pp. 261–272. DOI: 10.1038/s41592-019-0686-2.

[23] Marco Cuturi. “Sinkhorn Distances: Lightspeed Computation of Optimal Transport”. In: Advances in Neural Information Processing Systems. Vol. 26. Curran Associates, Inc., 2013, pp. 2292–2300. URL: https://proceedings.neurips.cc/paper/2013/hash/af21d0c97db2e27e13572cbf59eb343d-Abstract.html.

[24] Lénaïc Chizat et al. “Scaling algorithms for unbalanced optimal transport problems”. In: Mathematics of Computation 87.314 (2018), pp. 2563–2609. DOI: 10.1090/mcom/3303. URL: https://www.ams.org/mcom/2018-87-314/S0025-5718-2018-03303-8/.

[25] Marco Cuturi et al. “Optimal Transport Tools (OTT): A JAX Toolbox for all things Wasserstein”. In: arXiv preprint arXiv:2201.12324 (2022).

[26] 10x Genomics. High resolution mapping of the breast cancer tumor microenvironment using integrated single cell, spatial and in situ analysis of FFPE tissue (Xenium preview data, human breast). Dataset. In Situ Sample 2; Xenium Onboard Analysis 1.4. Accessed 23 October 2026. URL: https://www.10xgenomics.com/products/xenium-in-situ/preview-dataset-human-breast.

[27] Sunny Z. Wu et al. “A single-cell and spatially resolved atlas of human breast cancers”. In: Nature Genetics 53.9 (2021), pp. 1334–1347. DOI: 10.1038/s41588-021-00911-1. URL: https://www.nature.com/articles/s41588-021-00911-1.

[28] 10x Genomics. Preview Data: FFPE Human Lung Cancer with Xenium Multimodal Cell Segmentation. Dataset. Xenium Human Multi-Tissue and Cancer Panel, 377 genes; Xenium Onboard Analysis 2.0.0. Accessed 23 October 2026. URL: https://www.10xgenomics.com/datasets/preview-data-ffpe-human-lung-cancer-with-xenium-multimodal-cell-segmentation-1-standard.

[29] Nayoung Kim, Hong Kwan Kim, Kyungjong Lee, et al. “Single-cell RNA sequencing demonstrates the molecular and cellular reprogramming of metastatic lung adenocarcinoma”. In: Nature Communications 11.1 (2020), p. 2285. DOI: 10.1038/s41467-020-16164-1. URL: https://www.nature.com/articles/s41467-020-16164-1.

[30] Austin Hartman and Rahul Satija. “Comparative analysis of multiplexed in situ gene expression profiling technologies”. In: eLife (2024). Reviewed preprint. DOI: 10.7554/eLife.96949.

[31] Marius Pachitariu, Michael Rariden, and Carsen Stringer. “Cellpose-SAM: superhuman generalization for cellular segmentation”. In: bioRxiv (2025). Preprint. DOI: 10.1101/2025.04.28.651001.

[32] Leland McInnes et al. “UMAP: Uniform Manifold Approximation and Projection”. In: Journal of Open Source Software 3.29 (2018), p. 861. DOI: 10.21105/joss.00861. URL: https://joss.theoj.org/papers/10.21105/joss.00861.

[33] Peter J. Rousseeuw. “Silhouettes: A graphical aid to the interpretation and validation of cluster analysis”. In: Journal of Computational and Applied Mathematics 20 (1987), pp. 53–65. DOI: 10.1016/0377-0427(87)90125-7. URL: https://www.sciencedirect.com/science/article/pii/0377042787901257.

[34] Ilya Korsunsky et al. “Fast, sensitive and accurate integration of single-cell data with Harmony”. In: Nature Methods 16.12 (2019), pp. 1289–1296. DOI: 10.1038/s41592-019-0619-0. URL: https://www.nature.com/articles/s41592-019-0619-0.

[35] Yasin Şenbabaoğlu et al. “Tumor immune microenvironment characterization in clear cell renal cell carcinoma identifies prognostic and immunotherapeutically relevant messenger RNA signatures”. In: Genome Biology 17.1 (2016), p. 231. DOI: 10.1186/s13059-016-1092-z.

[36] Catherine S. Grasso et al. “Conserved interferon-γ signaling drives clinical response to immune check-point blockade therapy in melanoma”. In: Cancer Cell 38.4 (2020), 500–515.e3. DOI: 10.1016/j.ccell.2020.08.005.

[37] Angela M. Gocher, Creg J. Workman, and Dario A. A. Vignali. “Interferon-γ: teammate or opponent in the tumour microenvironment?” In: Nature Reviews Immunology 22.3 (2022), pp. 158–172. DOI: 10.1038/s41577-021-00566-3.

[38] Karthik Dhatchinamoorthy, Jeff D. Colbert, and Kenneth L. Rock. “Cancer immune evasion through loss of MHC class I antigen presentation”. In: Frontiers in Immunology 12 (2021), p. 636568. DOI: 10.3389/fimmu.2021.636568.

[39] Mark Ayers et al. “IFN-γ–related mRNA profile predicts clinical response to PD-1 blockade”. In: Journal of Clinical Investigation 127.8 (2017), pp. 2930–2940. DOI: 10.1172/JCI91190.

[40] Jesse M. Zaretsky et al. “Mutations associated with acquired resistance to PD-1 blockade in melanoma”. In: New England Journal of Medicine 375.9 (2016), pp. 819–829. DOI: 10.1056/NEJMoa1604958.

[41] jinworks. SpatialCellChat: CellChat for spatial transcriptomics. GitHub repository. Accessed 21 October 2026. URL: https://github.com/jinworks/SpatialCellChat.

[42] Suoqin Jin, Maksim V. Plikus, and Qing Nie. “CellChat for systematic analysis of cell–cell communication from single-cell transcriptomics”. In: Nature Protocols 20.1 (2025). First published online 16 September 2024, pp. 180–219. DOI: 10.1038/s41596-024-01045-4. URL: https://www.nature.com/articles/s41596-024-01045-4.

[43] Mariia Bilous et al. “Resolving sensitivity, specificity and signal contamination in Xenium spatial transcriptomics”. In: Nature Methods 23.6 (2026), pp. 1152–1162. DOI: 10.1038/s41592-026-03089-8.

[44] Can Ergen and Nir Yosef. “ResolVI –addressing noise and bias in spatial transcriptomics”. In: bioRxiv (2025). DOI: 10.1101/2025.01.20.634005.

[45] Y. Yang et al. “MisTIC: Missegmented Transcript Inference Correction for Improved Spatial Transcriptomics Analysis”. In: bioRxiv (2025). DOI: 10.64898/2025.12.11.693759.

[46] J. Wang et al. “Semi-metric optimal transport enables robust spatial multi-omics integration”. In: Research Square (2026). Preprint, version 1. DOI: 10.21203/rs.3.rs-10505162/v1.

