## Supplementary Notes 1-3, Tables S1-S3 and Figures S1-S2 for "Unified cell segmentation for spatial transcriptomics via transcript reassignment"

#### 1 Supplementary Notes

##### 1.1 Supplementary Note 1: Cell-type labels used in cell OT

The structure term of cell OT compares cell-type labels only within each domain, so the spatial and reference labels do not need to have the same names. For Xenium breast S1, the 10x author annotations [1] and the minor cell types of the breast cancer atlas of Wu et al. [2] were used as given. For Xenium breast S2 [3] and Visium HD colon [4], the spatial and reference labels come from the same annotation scheme of the respective study and were used directly. For CosMx lung [5] and Xenium lung [6], the spatial annotations are much finer than the broad cell types of the tumor-lung reference GSE131907 [7], so both domains were mapped to common broad groups before cell OT: eight groups for CosMx lung (Table S1) and seven for Xenium lung (Table S2). The broad groups were used only in the structure term of cell OT; all evaluations use the finer spatial labels.

Xenium lung has no released cell-type annotation. We annotated the 24 graph-based clusters of the 10x Xenium Onboard Analysis output [6] manually from the official cluster differential-expression markers, giving 15 fine labels (Table S3). In GSE131907, T lymphocytes and NK cells were merged into T/NK because the Xenium clustering has a single *CD3D*/*CD3E*/*TRAC*/*NKG7* lymphocyte cluster, and smooth muscle and pericyte clusters were assigned to Fibroblast, consistent with the fibroblast group of GSE131907, which contains smooth muscle cells, pericytes and myofibroblasts. Five historical GSE131907 gene symbols were renamed to their current symbols (*PTRF*→*CAVIN1*, *SDPR*→*CAVIN2*, *HRSP12*→*RIDA*, *C8orf4*→*TCIM*, *FAM46C*→*TENT5C*), giving 374 shared genes.

##### 1.2 Supplementary Note 2: Gene pair F1 score

We assessed transcript assignment with pairs of genes whose co-detection within a cell is either expected or suspicious, defined from the subcellular location of transcripts in the Original segmentation. Transcripts inside the nucleus are almost certainly from the cell itself, whereas cytoplasmic transcripts may come from neighbors. For a gene pair $(a, b)$, we therefore compared how often nuclear transcripts of gene $a$ co-occur with cytoplasmic transcripts of gene $b$ in the same cell (own) versus in an adjacent cell (neighbor). Adjacent cells were the six nearest cells within $25\,\mu$m, restricted to neighbors of a different cell type, and only cells with at least five nuclear and five cytoplasmic counts were used. Both values were computed from per-cell expression fractions and summed over all adjacent cell pairs and over both orders of the two genes. The enrichment of a pair was the $\log_2$ ratio of own to neighbor co-occurrence, centered by the median ratio of each gene to remove gene-specific differences between nucleus and cytoplasm, together with a $z$-like statistic, that is, the own–neighbor difference divided by its root-mean-square variation across adjacent cell pairs, which removes pairs driven by a few cells. "Intrinsic pairs" co-occur mainly within a cell ($\log_2$ ratio $\geq 1.5$, $z \geq 5$), and their co-detection in one cell is expected. "Suspicious pairs" co-occur mainly between adjacent cells of different types ($\log_2$ ratio $\leq -0.75$, $z \leq -5$; $\log_2$ ratio $\leq -0.5$ for Xenium breast S1), so their co-detection in one cell most likely reflects transcripts assigned to the wrong cell. This yielded 249 intrinsic and 213 suspicious pairs in Xenium breast S1, 595 and 381 in Xenium breast S2, 149 and 244 in CosMx lung, and 675 and 307 in Xenium lung.

For each method, a pair was co-detected in a cell if both genes had at least one count. Precision measures how specific co-detection is to intrinsic pairs: we averaged the number of co-detecting cells over intrinsic pairs

($\bar{n}_{\text{int}}$) and over suspicious pairs ($\bar{n}_{\text{sus}}$), and defined precision as $\bar{n}_{\text{int}}/(\bar{n}_{\text{int}}+\bar{n}_{\text{sus}})$, so that the two groups are weighted equally regardless of their sizes. Recall measures how many expected co-detections are recovered. Because methods produce different numbers of cells, we compared them on a common square grid whose side equals the typical cell size of the dataset (the square root of the median area per cell, 9.6–12.4 $\mu$m). An intrinsic pair was found in a grid square if any cell of the method with its centroid in that square co-detected the pair. Recall was the number of (square, intrinsic pair) combinations found by the method divided by the number found by any method, averaged over four grid offsets (shifts of half a square in $x$, $y$ or both). The F1 score is the harmonic mean of precision and recall.

##### 1.3 Supplementary Note 3: Differential expression between tumor cells near and far from T cells

We used the CosMx lung dataset and the original cell-type annotation for both Original and Tracel, so that the two methods were compared on the same cells with the same labels and centroids. Tumor cells were the cells of the five tumor clusters ($n$ = 19,186), and T cells were the CD4 and CD8 naive and memory T cells and regulatory T cells. For each tumor cell, we counted the T cells within 50 $\mu$m and computed the distance to the nearest T cell. Tumor cells with at least three T cells within 50 $\mu$m were defined as near ($n$ = 625), tumor cells whose nearest T cell was more than 100 $\mu$m away as far ($n$ = 7,857), and the remaining tumor cells were excluded from the comparison. Counts of each cell were scaled to the median total count per cell, the same target value for both methods. We tested each gene between near and far cells with the two-sided Wilcoxon rank-sum test on log-transformed normalized counts, separately for each method, corrected $P$-values by the Benjamini–Hochberg procedure [8], and computed the $\log_2$ fold change from the mean normalized counts with a pseudocount of 0.05.

We used two published gene sets as the expected IFN-$\gamma$ response of tumor cells. The antigen-presentation machinery (APM) set is the core MHC class I pathway, which is expressed by tumor cells and induced by IFN-$\gamma$. It contains *HLA-A*, *HLA-B*, *HLA-C*, *B2M*, *TAP1* and *TAP2* (*TAPBP* is not on the panel) [9]. The IFN-$\gamma$-induced set was measured directly in tumor cells: it contains the 549 genes significantly up-regulated after IFN-$\gamma$ treatment of melanoma cell lines [10], 50 of which are on the panel. As a contamination control, we used 15 canonical T-cell markers on the panel (*CCL5*, *CD2*, *CD3D*, *CD3E*, *CD3G*, *CD8A*, *CD8B*, *CD28*, *GZMA*, *GZMB*, *GZMK*, *IL7R*, *ITK*, *NKG7*, *PRF1*), all of which are marker genes of lung T cells in the human lung cell atlas [11]; these genes are not expected to differ between the two groups of tumor cells. Gene set enrichment was tested by preranked GSEA [12] with gseapy [13] (1,000 permutations), with genes ranked by the sign of the $\log_2$ fold change times $-\log_{10} p$, against the Hallmark collection [14] together with the gene sets above, and FDR was computed over all tested sets. To show how expression changes with distance, all tumor cells were grouped by the distance to the nearest T cell (0–15, 15–30, 30–50, 50–100, 100–200 and $> 200\,\mu$m), and for each gene set we report the mean and 95% confidence interval of the summed normalized counts of its genes per cell.

#### 2 Supplementary Tables

**Table S1: Cell-type label mapping for CosMx lung.** Official NanoString annotations of Lung5 Rep2 [5] and the cell types of the tumor-lung cells of GSE131907 [7] were mapped to eight broad groups for the structure term of cell OT. Numbers of cells are in parentheses (105,800 spatial cells; 45,149 reference cells).

| Broad group | CosMx labels (cells) | GSE131907 label (cells) |
|---|---|---|
| Epithelial | tumor 5 (17,982), tumor 12 (1,175), tumor 9 (19), tumor 13 (6), tumor 6 (4), epithelial (4,059) | Epithelial cells (7,270) |
| Myeloid | neutrophil (13,377), macrophage (6,857), mDC (4,325), pDC (2,175), monocyte (1,794) | Myeloid cells (8,794) |
| T | T CD4 naive (9,576), T CD8 naive (1,935), T CD4 memory (193), Treg (64), T CD8 memory (28) | T lymphocytes (18,587) |
| B | plasmablast (8,675), B-cell (8,273) | B lymphocytes (5,312) |
| Fibroblast | fibroblast (12,670) | Fibroblasts (1,718) |
| Endothelial | endothelial (8,075) | Endothelial cells (655) |
| Mast | mast (2,729) | MAST cells (1,809) |
| NK | NK (1,809) | NK cells (1,004) |

**Table S2: Cell-type label mapping for Xenium lung.** Spatial labels from our annotation of the Xenium graph-based clusters [6] (Table S3) and the cell types of the tumor-lung cells of GSE131907 [7] were mapped to seven broad groups for the structure term of cell OT. Numbers of cells are in parentheses (49,786 spatial cells; 45,147 reference cells).

| Broad group | Xenium lung labels (cells) | GSE131907 label (cells) |
|---|---|---|
| Epithelial | Tumor epithelial (24,052), Respiratory ciliated epithelial (1,807), Alveolar epithelial (331), Cycling tumor epithelial (245) | Epithelial cells (7,270) |
| Myeloid | Macrophage/monocyte (3,241), Dendritic/myeloid (1,116), Macrophage with epithelial spillover (1,018), Macrophage (299) | Myeloid cells (8,794) |
| T/NK | T/NK lymphocytes (7,014) | T lymphocytes (18,586), NK cells (1,004) |
| B | Plasma/B lymphocyte (1,784), B lymphocyte (877) | B lymphocytes (5,311) |
| Fibroblast | Fibroblast (4,477), Smooth muscle/pericyte (747) | Fibroblasts (1,718) |
| Endothelial | Endothelial (2,226) | Endothelial cells (655) |
| Mast | Mast (552) | MAST cells (1,809) |

**Table S3: Cell-type annotation of Xenium lung.** The 24 graph-based clusters of the Xenium Onboard Analysis output [6] were annotated manually from the official cluster differential-expression markers listed, giving 15 labels.

| Label | Clusters | Marker genes |
|---|---|---|
| Tumor epithelial | 1, 2, 3, 6 | *EPCAM*, *KRT7*, *GPRC5A*, *CFTR*, *CYP2B6*, *CYP2A7*, *KLK11*, *TCIM*, *MALL*, *CAPN8*, *CYP4B1*, *SFTA2*, *COL17A1*, *MYC*, *MET*, *EHF* |
| Respiratory ciliated epithelial | 10, 22 | *FOXJ1*, *CCDC39*, *DNAAF1*, *SNTN*, *CFAP53*, *C20orf85* |
| Alveolar epithelial | 15 | *AGER*, *LAMP3*, *CAV1*, *PDPN*, *RTKN2* |
| Cycling tumor epithelial | 17 | *MKI67*, *TOP2A*, *CENPF*, *CCNB2*, *EPCAM*, *KRT7* |
| Macrophage/monocyte | 7 | *CD163*, *MRC1*, *MS4A4A*, *CD14*, *FCGR3A*, *CD68* |
| Dendritic/myeloid | 13 | *CD1C*, *CD1E*, *FCER1A*, *CLEC10A*, *IRF8*, *FCN1* |
| Macrophage with epithelial spillover | 8 | *MARCO*, *MCEMP1*, *FCGR3A*, *CD68*, *AIF1*, *MRC1* |
| Macrophage | 16 | *MARCO*, *MCEMP1*, *MRC1*, *FCGR3A*, *CD68*, *CD163* |
| T/NK lymphocytes | 4 | *CD2*, *CD3D*, *CD3E*, *TRAC*, *IL7R*, *CD8A*, *NKG7* |
| Plasma/B lymphocyte | 12 | *MZB1*, *CD79A*, *TNFRSF17*, *PRDM1*, *SLAMF7*, *CD19* |
| B lymphocyte | 18, 19 | *MS4A1*, *CD19*, *CD79A*, *BANK1*, *SPIB* |
| Fibroblast | 5 | *PDGFRA*, *COL5A2*, *SFRP2*, *SFRP4*, *FBN1*, *LTBP2* |
| Smooth muscle/pericyte | 14, 20 | *MYH11*, *CNN1*, *ACTA2*, *DES*, *MYLK*, *OGN* |
| Endothelial | 9, 11 | *VWF*, *ACKR1*, *CLEC14A*, *ADGRL4*, *RAMP2*, *EGFL7*, *CD34*, *KCNK3*, *EDNRB* |
| Mast | 21, 23, 24 | *CPA3*, *MS4A2*, *KIT*, *IL1RL1*, *SLC18A2*, *HPGDS* |

### 3 Supplementary Figures

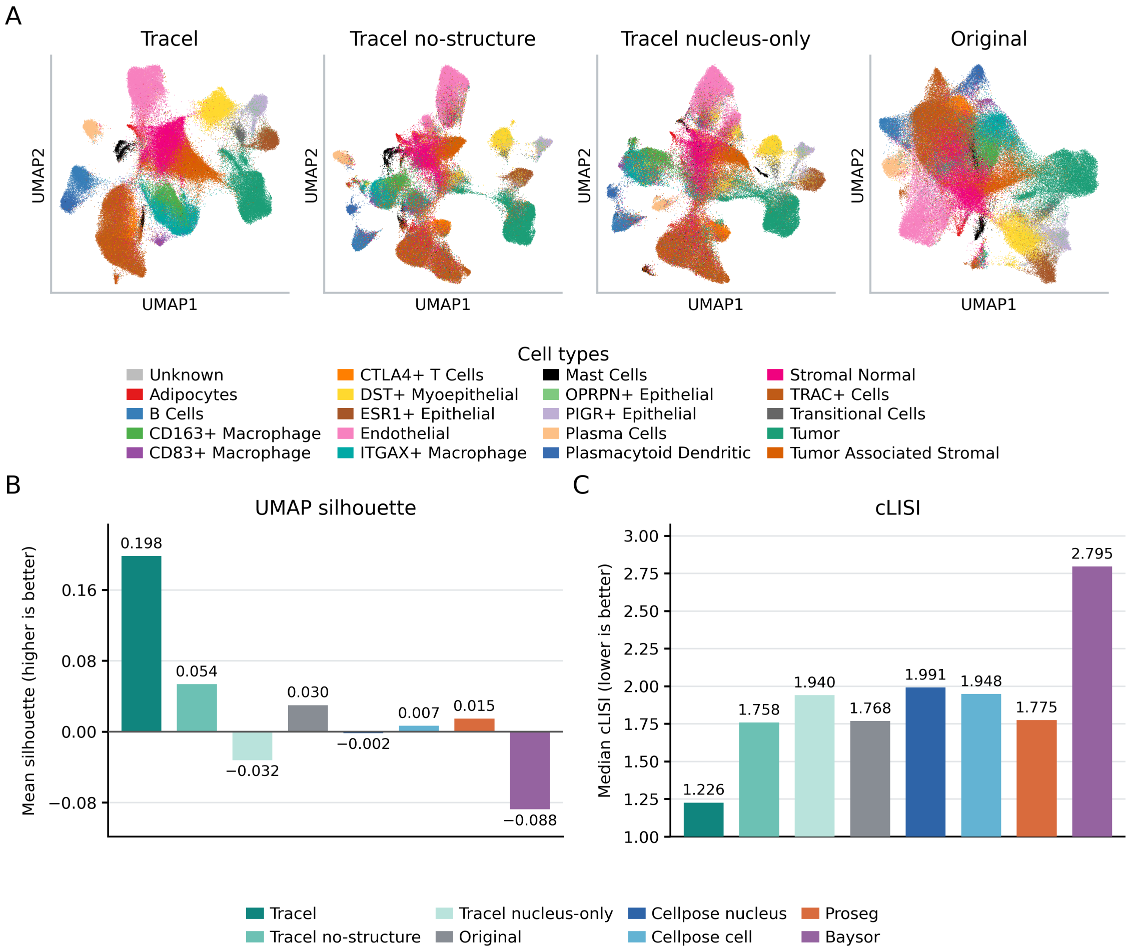


**Figure S1: Effect of the cell-type structure term and of nucleus-only input in cell OT, Xenium breast S1. A**, UMAP embeddings colored by the published cell-type annotations, of Tracel, Tracel without the cell-type structure term (Tracel no-structure), Tracel no-structure with cell OT started from nucleus segmentation, that is, using only the transcripts located inside nuclei in cell OT (Tracel nucleus-only); and the Original segmentation. All Tracel variants use the same transcript OT on all transcripts. Tracel and Original are the embeddings of Fig. 2A. **B**, Mean UMAP silhouette and **C**, median cell-type LISI (cLISI) from 30 nearest neighbors in each method's 30-component PCA embedding, computed as in Fig. 2B and C from 5,000 sampled cells per method. Values of the other methods are those of Fig. 2B and C. Higher silhouette and lower cLISI indicate better separation of cell types.

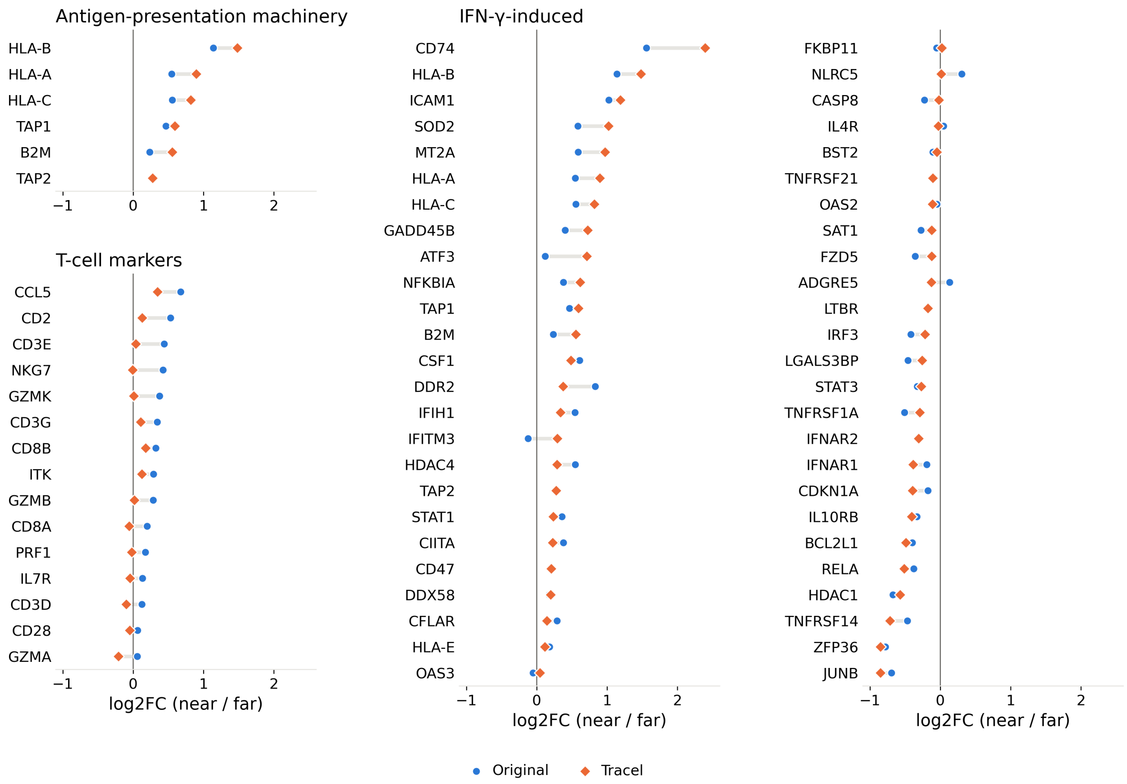


**Figure S2: Per-gene expression differences between tumor cells near and far from T cells.** $\text{Log}_2$ fold change of mean normalized counts between near tumor cells (at least three T cells within 50 $\mu$m; $n = 625$) and far tumor cells (nearest T cell more than 100 $\mu$m away; $n = 7{,}857$) in the CosMx lung dataset, for Original (circles) and Tracel (diamonds). Left, antigen-presentation machinery genes (top) and T-cell markers (bottom), which are not expected to differ between tumor cells; middle and right, IFN-$\gamma$-induced genes.

### Supplementary References


[1] 10x Genomics. *High resolution mapping of the breast cancer tumor microenvironment using integrated single cell, spatial and in situ analysis of FFPE tissue (Xenium preview data, human breast)*. Dataset. In Situ Sample 2; Xenium Onboard Analysis 1.4. Accessed 23 October 2026. URL: https://www.10xgenomics.com/products/xenium-in-situ/preview-dataset-human-breast.

[2] Sunny Z. Wu et al. "A single-cell and spatially resolved atlas of human breast cancers". In: *Nature Genetics* 53.9 (2021), pp. 1334–1347. DOI: 10.1038/s41588-021-00911-1. URL: https://www.nature.com/articles/s41588-021-00911-1.

[3] Amanda Janesick et al. "High resolution mapping of the tumor microenvironment using integrated single-cell, spatial and in situ analysis". In: *Nature Communications* 14.1 (2023), p. 8353. DOI: 10.1038/s41467-023-43458-x. URL: https://www.nature.com/articles/s41467-023-43458-x.

[4] Michelli Faria de Oliveira et al. "High-definition spatial transcriptomic profiling of immune cell populations in colorectal cancer". In: *Nature Genetics* 57.6 (2025), pp. 1512–1523. DOI: 10.1038/s41588-025-02193-3. URL: https://www.nature.com/articles/s41588-025-02193-3.

[5] Shanshan He, Ruchir Bhatt, Carl Brown, et al. "High-plex imaging of RNA and proteins at subcellular resolution in fixed tissue by spatial molecular imaging". In: *Nature Biotechnology* 40.12 (2022), pp. 1794–1806. DOI: 10.1038/s41587-022-01483-z. URL: https://www.nature.com/articles/s41587-022-01483-z.

[6] 10x Genomics. *Preview Data: FFPE Human Lung Cancer with Xenium Multimodal Cell Segmentation*. Dataset. Xenium Human Multi-Tissue and Cancer Panel, 377 genes; Xenium Onboard Analysis 2.0.0. Accessed 23 October 2026. URL: https://www.10xgenomics.com/datasets/preview-data-ffpe-human-lung-cancer-with-xenium-multimodal-cell-segmentation-1-standard.

[7] Nayoung Kim, Hong Kwan Kim, Kyungjong Lee, et al. "Single-cell RNA sequencing demonstrates the molecular and cellular reprogramming of metastatic lung adenocarcinoma". In: *Nature Communications* 11.1 (2020), p. 2285. DOI: 10.1038/s41467-020-16164-1. URL: https://www.nature.com/articles/s41467-020-16164-1.

[8] Yoav Benjamini and Yosef Hochberg. "Controlling the false discovery rate: a practical and powerful approach to multiple testing". In: *Journal of the Royal Statistical Society: Series B (Methodological)* 57.1 (1995), pp. 289–300. DOI: 10.1111/j.2517-6161.1995.tb02031.x.

[9] Yasin Şenbabaoğlu et al. "Tumor immune microenvironment characterization in clear cell renal cell carcinoma identifies prognostic and immunotherapeutically relevant messenger RNA signatures". In: *Genome Biology* 17.1 (2016), p. 231. DOI: 10.1186/s13059-016-1092-z.

[10] Catherine S. Grasso et al. "Conserved interferon-$\gamma$ signaling drives clinical response to immune checkpoint blockade therapy in melanoma". In: *Cancer Cell* 38.4 (2020), 500–515.e3. DOI: 10.1016/j.ccell.2020.08.005.

[11] Kyle J. Travaglini et al. "A molecular cell atlas of the human lung from single-cell RNA sequencing". In: *Nature* 587.7835 (2020), pp. 619–625. DOI: 10.1038/s41586-020-2922-4.

[12] Aravind Subramanian et al. "Gene set enrichment analysis: a knowledge-based approach for interpreting genome-wide expression profiles". In: *Proceedings of the National Academy of Sciences* 102.43 (2005), pp. 15545–15550. DOI: 10.1073/pnas.0506580102.

[13] Zhuoqing Fang, Xinyuan Liu, and Gary Peltz. "GSEApy: a comprehensive package for performing gene set enrichment analysis in Python". In: *Bioinformatics* 39.1 (2023), btac757. DOI: 10.1093/bioinformatics/btac757.

[14] Arthur Liberzon et al. "The Molecular Signatures Database hallmark gene set collection". In: *Cell Systems* 1.6 (2015), pp. 417–425. DOI: 10.1016/j.cels.2015.12.004.
